# Alignment-Free Microhaplotype Genotyping for GT-seq (Genotyping-in-Thousands by Sequencing) Using a Diploid Abundance Model

**DOI:** 10.64898/2026.04.01.715880

**Authors:** Nathan R Campbell, Amanda R Campbell, Shannon K Blair, Amanda J Finger

## Abstract

GT-seq (Genotyping-in-Thousands by Sequencing) is widely used for high-throughput amplicon genotyping, but most analytical pipelines focus on single SNPs or rely on alignment-based variant calling. Here we present an alignment-free approach for microhaplotype genotyping that leverages the high read depth and low error rates typical of paired-end Illumina and Element sequencing. The pipeline first identifies primer-bounded reads and resolves paired-end sequences into complete amplicon sequences. Within each sample and locus, unique sequences are ranked by read abundance and the top one or two sequences are retained as candidate diploid alleles. These alleles are aggregated across samples to construct a catalog of unique haplotypes for each locus. In a second pass, reads are assigned to catalog haplotypes by exact sequence matching to produce diploid genotypes. Finally, catalog haplotype sequences are positionally compared to identify phased SNP and collapsed indel variation, generating compact microhaplotype representations suitable for population genetic analysis. This approach enables robust, alignment-free microhaplotype inference directly from high-depth amplicon sequencing data.

## Introduction

Targeted amplicon sequencing has become a widely adopted approach for high-throughput genotyping in ecological, conservation, and breeding studies. Among these methods, Genotyping-in-Thousands by sequencing (GT-seq) has emerged as an efficient and cost-effective strategy for genotyping large numbers of individuals at hundreds of loci simultaneously (Campbell et al. 2015). GT-seq uses highly multiplexed PCR amplification of targeted genomic regions followed by next-generation sequencing, allowing thousands of individuals to be genotyped in a single sequencing run. The method has been widely applied in studies of population structure, mixed-stock analysis, hybridization, and genetic monitoring of natural populations.

Since its introduction, GT-seq panels have been developed for a broad range of species and research applications. The method is particularly useful in systems where large numbers of individuals must be genotyped across moderately sized marker panels. For example, GT-seq panels have been implemented for genetic monitoring and population structure analyses in wildlife and fisheries systems where large sample sizes are required to achieve sufficient statistical power (Schmidt et al. 2020; Hayward et al. 2022). Because the method relies on targeted amplification rather than genome-wide sequencing, GT-seq provides a scalable and cost-effective alternative for studies requiring genotyping of thousands of individuals.

An important application of GT-seq and other targeted sequencing methods is parentage and kinship analysis. High-throughput SNP panels generated using GT-seq have been used to assign parentage, reconstruct pedigrees, and estimate relatedness in natural populations (Campbell et al. 2015; Hargrove et al. 2021; Schmidt et al. 2020). The ability to genotype large numbers of individuals at relatively low cost makes GT-seq particularly well suited for kinship-based studies in species with large population sizes or complex breeding systems.

Most analytical pipelines developed for GT-seq data treat each locus as a single SNP marker or rely on alignment-based variant calling approaches in which sequencing reads are mapped to a reference prior to variant discovery (Campbell et al. 2015; Baetscher et al. 2018). While effective, these approaches do not fully exploit the structure of targeted amplicon sequencing data. Because each amplicon typically spans a short genomic region, multiple polymorphic sites may occur within the same amplicon and can be observed simultaneously on individual sequencing reads. These combinations of linked polymorphisms form haplotypes that may contain substantially more information than individual SNP markers.

Traditional haplotype inference approaches often rely on statistical phasing methods that reconstruct haplotypes from genotype data (Browning and Browning 2011). In contrast, targeted amplicon sequencing frequently produces reads that span the entire amplified region, allowing haplotypes to be observed directly from sequencing reads without statistical inference. In such cases, alignment-based SNP pipelines introduce unnecessary analytical complexity by first reducing reads to independent SNP calls and subsequently reconstructing haplotypes from these variants.

Microhaplotypes, defined as short genomic regions containing two or more closely spaced polymorphisms observed on the same DNA fragment, have emerged as powerful genetic markers for applications including ancestry inference, individual identification, and kinship analysis (Kidd et al. 2014; Pakstis et al. 2021). Because multiple SNPs are observed together on the same sequencing read, microhaplotypes often produce multi-allelic markers with higher heterozygosity and greater discriminatory power than single SNP loci.

Several studies have demonstrated that microhaplotypes can substantially improve the statistical power of relatedness analyses. Because each locus may produce multiple haplotype alleles rather than two SNP states, microhaplotypes increase the effective information content of genetic markers and reduce false-positive rates in kinship inference (Anderson et al. 2018; Kidd et al. 2014; Pakstis et al. 2021). Comparative studies have shown that microhaplotype marker panels can outperform equivalent SNP panels in distinguishing closely related individuals and resolving complex pedigree relationships (Baetscher et al. 2018; Tomas et al. 2024).

Targeted amplicon sequencing approaches such as GT-seq are therefore particularly well suited for microhaplotype genotyping. Sequencing reads frequently span the entire amplified region, allowing haplotypes to be reconstructed directly from sequencing reads rather than inferred statistically from independent SNP calls. Furthermore, the high sequencing depth typical of GT-seq experiments provides a strong signal for distinguishing true allele sequences from sequencing errors based on read abundance.

Here we describe an alignment-free analytical framework for microhaplotype genotyping from GT-seq amplicon sequencing data. The method resolves paired-end reads into primer-bounded amplicon sequences, identifies candidate allele sequences within each sample using a diploid abundance model, and constructs a catalog of unique haplotypes across all samples. In a second pass, sequencing reads are assigned to catalog haplotypes by exact sequence matching to produce diploid genotype calls. Catalog haplotypes are then positionally compared to identify polymorphic sites and generate phased microhaplotype representations suitable for downstream population genetic and kinship analyses. Because many GT-seq amplicon panels already include loci that span multiple polymorphic sites, existing GT-seq panels may be analyzed using this pipeline without modification to laboratory protocols.

## Materials and Methods

The analysis described here is implemented in a Python-based software script developed for processing paired-end amplicon sequencing data generated by GT-seq panels. The script accepts paired-end FASTQ files and a table of primer sequences defining the targeted loci. The software performs four primary steps: (1) resolution of paired-end reads into primer-bounded amplicon sequences, (2) identification of candidate allele sequences and construction of a locus-specific allele catalog, (3) genotype inference using an abundance-based diploid model, and (4) extraction of phased microhaplotype representations from catalog haplotypes. The implementation is designed to operate directly on high-depth amplicon sequencing reads without requiring read alignment to a reference or use other conventional variant-calling pipelines. The software is designed to operate on standard paired-end FASTQ files such as those produced by Illumina or Element sequencing platforms.

### GT-seq Library Preparation and Sequencing

Genomic DNA samples from delta smelt (*Hypomesus transpacificus*) were genotyped using a GT-seq amplicon sequencing panel consisting of 410 targeted loci. The panel included loci selected from prior marker discovery efforts, including candidate microhaplotypes identified before panel development. GT-seq library preparation followed the general protocol described by Campbell et al. (2015), in which locus-specific primer pairs are multiplexed in a single PCR. However, the indexing and normalization steps were performed using Nate’s Plates normalization and tagging kits (GTseek LLC, Twin Falls, Idaho, USA). The pooled GT-seq library was sequenced using an Element AVITI sequencing instrument at the University of Minnesota Genomics Center (UMGC).

Sequencing was performed using paired-end (PE100) chemistry, producing paired FASTQ files for each individual sample.

To evaluate cross-platform reproducibility, the same pooled GT-seq library was subsequently sequenced on an Illumina NovaSeq X instrument using paired-end short-read chemistry. FASTQ files generated from the Illumina run were processed using the same analysis pipeline described above. For cross-platform comparison, genotype inference for the Illumina dataset was performed using the allele catalog constructed from the Element dataset, allowing direct comparison of genotype calls across sequencing platforms without re-generating locus-specific catalogs. This approach ensures that any differences observed reflect differences in sequencing data rather than differences in allele discovery or catalog construction.

### Ethics Statement

All fish sampling and handling were conducted by the Fish Conservation and Culture Laboratory (FCCL) in accordance with institutional and governmental guidelines. Fin clip samples were collected under approved animal care and use protocols (IACUC protocol number 23041).

### Example Dataset

To demonstrate the performance of the alignment-free microhaplotype pipeline, a subset of sequencing data from this GT-seq panel was used as an example dataset. The dataset consists of paired-end (PE100) FASTQ files from 96 delta smelt (*Hypomesus transpacificus*) individuals prepared with the 410-locus GT-seq panel. Delta smelt is a diploid species, making it appropriate for evaluation under the diploid abundance model implemented here. To reduce file size while preserving locus representation and genotype structure, sequencing reads were randomly subsampled to 150,000 read pairs per individual. The resulting dataset provides sufficient sequencing depth to demonstrate allele discovery, catalog construction, and genotype inference using the alignment-free microhaplotype pipeline.

The example dataset is publicly available at Zenodo: https://doi.org/10.5281/zenodo.19069551

### Primer-Bounded Paired-End Read Resolution

GT-seq libraries consist of PCR amplicons generated from locus-specific primer pairs and sequenced using paired-end short-read platforms. Because each locus is defined by known primer sequences, these primers provide natural anchors for identifying and reconstructing the targeted amplicon sequences.

The analysis pipeline scans paired-end FASTQ files to identify reads beginning with a forward primer sequence corresponding to a defined locus. Candidate reads are then evaluated using their paired read to confirm the presence of the corresponding reverse primer sequence. Only read pairs containing the expected primer combination are retained for downstream analysis.

For validated read pairs, the full amplicon sequence is reconstructed by merging the paired reads. In cases where the amplicon length is shorter than the sequencing read length, the entire amplicon may be contained within the first read. For longer amplicons, the overlapping regions of the paired reads are identified and merged using a fast ungapped overlap algorithm, allowing reconstruction of the full amplicon sequence.

All reconstructed sequences are normalized to a single orientation beginning with the forward primer and ending with the reverse complement of the reverse primer. The resulting primer-bounded amplicon sequences are written to locus-specific FASTQ files and represent the fundamental unit of analysis for subsequent allele discovery and genotyping.

### Allele Discovery and Catalog Construction

Allele discovery is performed using the primer-bounded amplicon sequences obtained from the read resolution stage. For each sample and locus, unique amplicon sequences are identified and their read counts tabulated. Because GT-seq data typically exhibit very high read depth per locus and Illumina and Element sequencers produce high-quality reads with low error rates, true allelic sequences occur at substantially higher frequencies than sequencing errors. Within each sample and locus, unique sequences are therefore ranked by read abundance. Under a diploid model, at most two alleles are expected per locus in each individual. Accordingly, the two most abundant sequences are retained as candidate alleles. Additional filtering is applied based on allele abundance ratios.

For example, when the most abundant sequence constitutes the vast majority of reads at a locus, the locus is classified as homozygous and only the dominant sequence is retained as the candidate allele. When two sequences occur at substantial frequencies, both are retained as candidate alleles representing a heterozygous genotype. Once all candidate alleles have been identified across all samples, the sequences are aggregated to construct a catalog of unique haplotypes for each locus.

Each distinct amplicon sequence is assigned a unique allele identifier. The resulting catalog represents the set of observed haplotypes across the dataset and serves as the reference for genotype calling in the second pass of the analysis.

### Genotype Inference Using a Diploid Abundance Model

Following construction of the allele catalog, the primer-bounded amplicon sequences are reanalyzed in a second pass to assign reads to catalog haplotypes. Each resolved sequence is compared against the allele catalog for the corresponding locus using exact sequence matching.

For each sample and locus, reads matching each catalog allele are counted. The two alleles with the highest read counts are retained as the candidate genotype for that sample. Genotype classification is then determined using the relative abundance of the second-most frequent allele.

Specifically, genotype classification is based on the proportion of reads corresponding to the second most abundant allele (A2). Samples with a second-allele proportion below a defined threshold (default = 10%) are classified as homozygous, whereas samples with substantial representation of two alleles (25% – 50%) are classified as heterozygous. Loci with insufficient read depth or ambiguous allele ratios are classified as low confidence or no-call. This abundance-based approach leverages the high sequencing depth typical of GT-seq assays to distinguish true allelic variation from sequencing noise.

### Microhaplotype Extraction and Phased SNP Representation

The allele catalog generated in the previous step consists of complete amplicon sequences representing observed haplotypes for each locus. Catalog haplotypes are positionally compared to identify polymorphic sites, allowing phased SNP and indel variation to be extracted. Because each allele sequence represents a contiguous DNA fragment observed in sequencing reads, the relative phase of polymorphisms within the locus is inherently preserved. Consequently, the resulting haplotypes represent phased combinations of SNPs and indels within each amplicon.

For each locus, the polymorphic sites are extracted from the catalog haplotypes to generate phased SNP representations of each allele. These phased haplotypes constitute microhaplotypes, defined as short genomic regions containing multiple linked polymorphisms that can be observed within a single sequencing read or reconstructed amplicon.

Microhaplotype genotypes are therefore represented as combinations of phased allele sequences rather than individual unlinked SNP calls. This representation captures the joint information from multiple polymorphic sites within each amplicon and can substantially increase the information content of loci used for kinship inference, parentage analysis, and population genetic applications.

### Software Implementation

The analysis workflow described above is implemented in the Python script gtseq_microhap_catalog_and_call.py, available through the GTseq GitHub repository: https://github.com/GTseq/gtseq_microhap

The software accepts paired-end FASTQ files and a table of locus-specific primer sequences as input and performs read resolution, allele catalog construction, genotype inference, and microhaplotype extraction in a single pipeline. The implementation is designed to operate directly on high-depth targeted amplicon sequencing data generated by GT-seq assays without requiring reference alignment or conventional variant-calling workflows.

## Results

### Sequencing Summary and Read Resolution Performance

The delta smelt demonstration dataset consisted of 96 individuals prepared using a 410-locus GT-seq panel. Across all samples, the sequencing run produced 234,935,633 raw read pairs, corresponding to an average of 2,447,246 raw read pairs per sample. Application of the primer-bounded read resolution step retained 164,209,863 primer-bounded read pairs, corresponding to an average of 1,710,519 primer-bounded read pairs per sample. Overall, 69.9% of raw read pairs were retained as primer-bounded reads suitable for downstream allele catalog construction and genotype inference. Among individual samples, raw read counts ranged from 1,255,265 to 3,270,346 read pairs, while primer-bounded read counts ranged from 166,594 to 2,602,617 read pairs. Most samples exhibited consistently high recovery of primer-bounded reads, although a small number of lower-performing samples were observed.

### Cross-Platform Genotype Concordance

To assess cross-platform reproducibility of the alignment-free microhaplotype pipeline, genotype calls generated from the Element dataset were compared to those obtained from the Illumina NovaSeq X sequencing run. Across 410 loci and 96 individuals, genotype concordance between platforms was 99.96% for loci successfully genotyped in both datasets (Figure S1). The majority of discrepancies were attributable to loci with low read depth or marginal genotype quality, resulting in no-call vs. genotype comparisons rather than conflicting genotype assignments. True conflicting genotype assignments between platforms were rare relative to comparisons involving no-calls.

Sample-level concordance between corresponding Element and Illumina samples exceeded 99.5% for all individuals, with most samples exhibiting greater than 99.9% genotype agreement (Table S1). Pairwise comparisons across all samples confirmed that each individual’s highest similarity was observed with its corresponding sample across platforms, with no evidence of sample misassignment or systematic platform bias (Figure S2).

### Public Example Dataset

To facilitate software testing and reproducibility, a trimmed example dataset was generated from the delta smelt sequencing run and deposited at Zenodo. This public dataset consists of paired-end FASTQ files subsampled to 150,000 read pairs per individual, together with the primer definition file required to run the pipeline. The example dataset is available at: https://doi.org/10.5281/zenodo.19069551

### Example Locus Demonstrating Microhaplotype Genotyping Model

To illustrate the method, locus NC_061065.1_5347094 was examined in detail. This locus contained multiple polymorphic positions, including both SNPs and a short indel variant. Because the allele sequences span the full amplicon, the phase of these linked polymorphisms was preserved directly in the catalog haplotypes allowing generation of phased SNP genotypes.

### Alignment-Free Microhaplotype Inference

Following read resolution, primer-bounded sequences were grouped by locus and unique amplicon sequences were ranked by read abundance within each sample. These candidate alleles were aggregated across samples to construct a catalog of observed haplotypes for each locus (Figure 3). In the second pass of the resolved FASTQ files, primer-bounded reads were assigned to catalog haplotypes by exact sequence matching, allowing diploid genotype inference based on allele abundance (Figure 4A).

**Figure 1.**
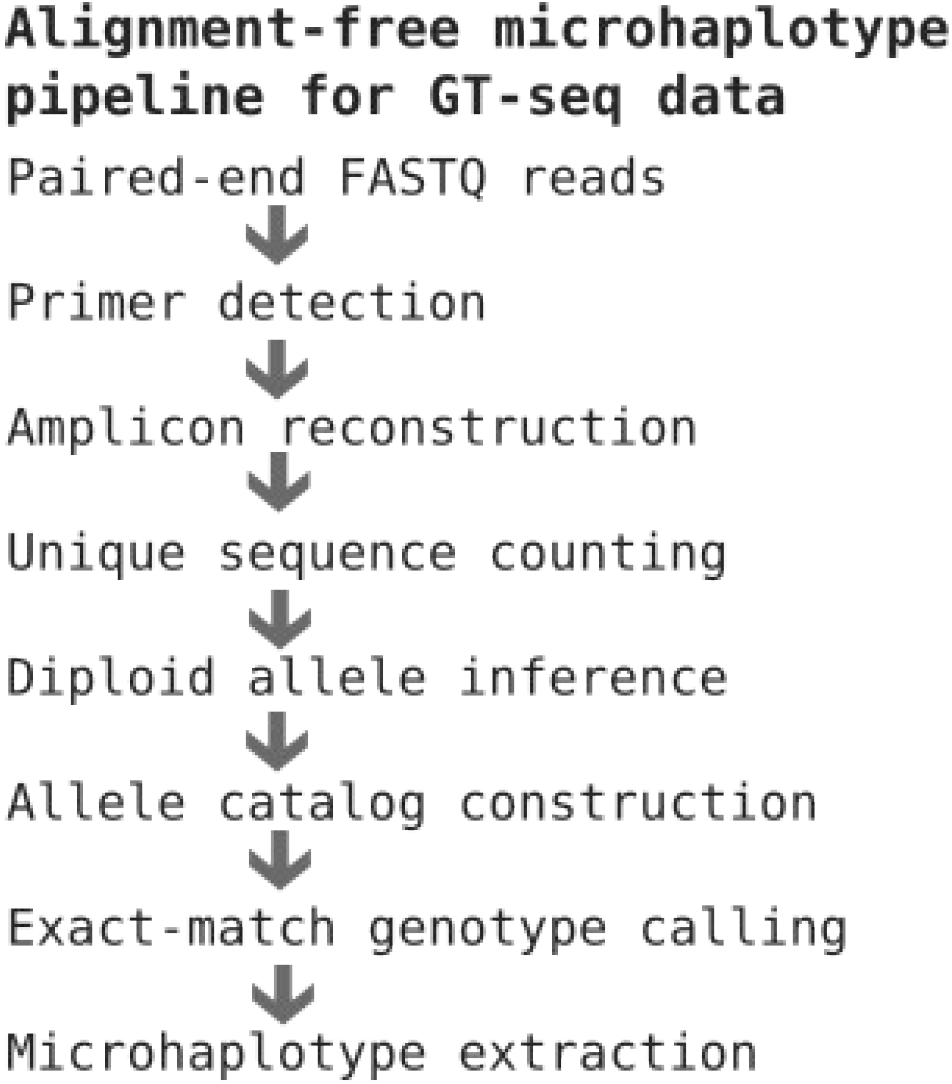
Alignment-free microhaplotype genotyping pipeline for GT-seq amplicon sequencing data. The analytical workflow implemented in the GT-seq microhaplotype pipeline processes paired-end FASTQ reads to infer diploid microhaplotype genotypes without reference alignment. Paired-end reads are first scanned to identify primer-bounded amplicons corresponding to targeted loci. Primer-bounded reads are resolved into complete amplicon sequences and grouped by locus. In the first pass of the analysis, unique amplicon sequences are identified within each sample and ranked by read abundance to infer candidate diploid alleles and construct a catalog of unique haplotypes for each locus. In the second pass, sequencing reads are assigned to catalog haplotypes by exact sequence matching. Allele counts are used to infer diploid genotypes using an abundance-based model, and polymorphic sites within catalog haplotypes are extracted to generate phased microhaplotype representations.

**Figure 2.**
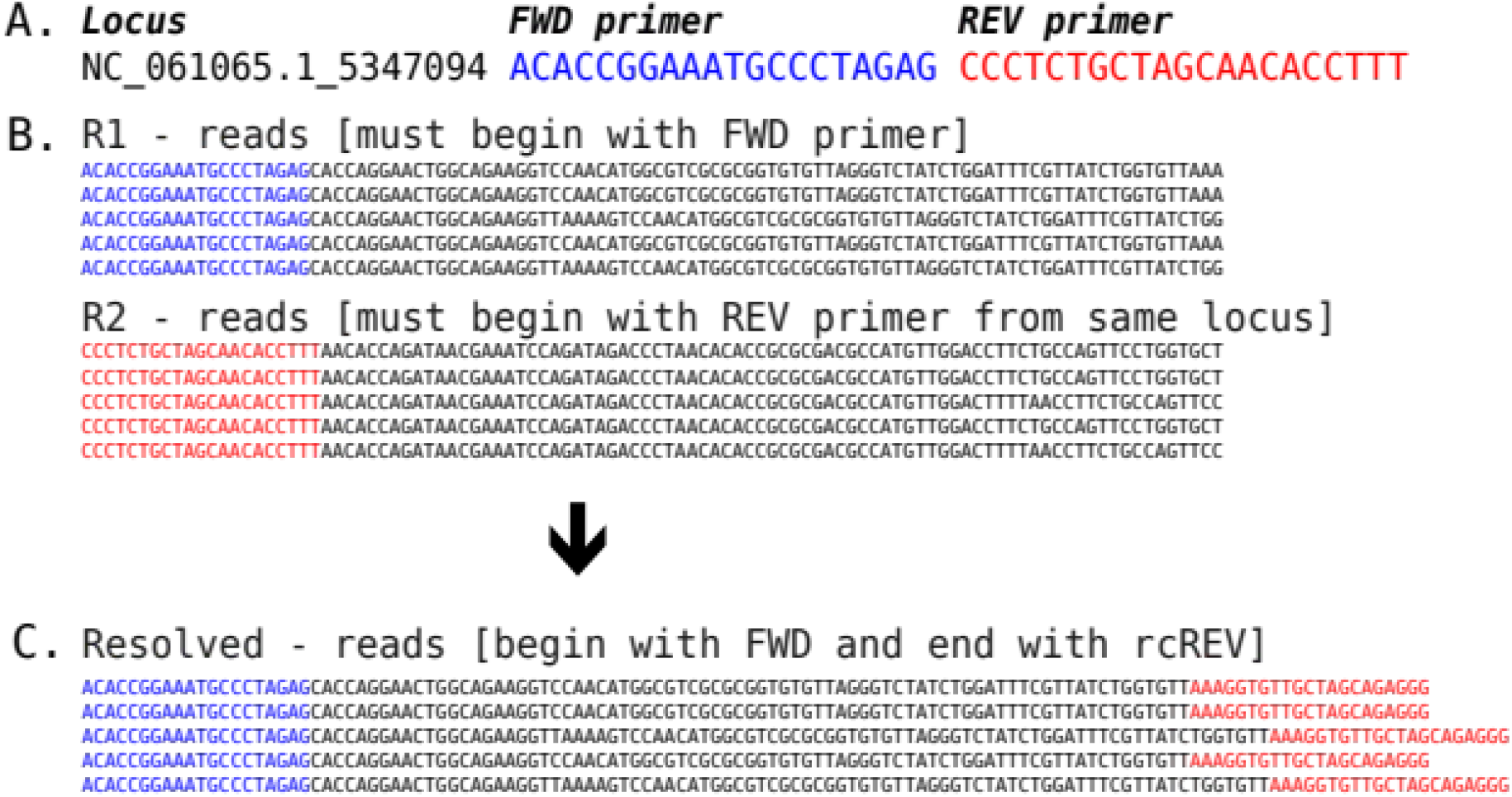
Reconstruction of primer-bounded amplicon sequences from paired-end reads. (A) Paired-end sequencing reads from GT-seq libraries contain locus-specific primer sequences that define the boundaries of each amplicon. Reads beginning with the forward primer are identified and paired with their corresponding reverse reads if the R2 read begins with the reverse primer string. (B) For validated read pairs, the full amplicon sequence is reconstructed by merging the paired reads. Overlapping regions of the paired reads are merged to reconstruct the full primer-bounded amplicon sequence, which begins with the forward primer and end with the reverse complement of the reverse primer.

**Figure 3.**
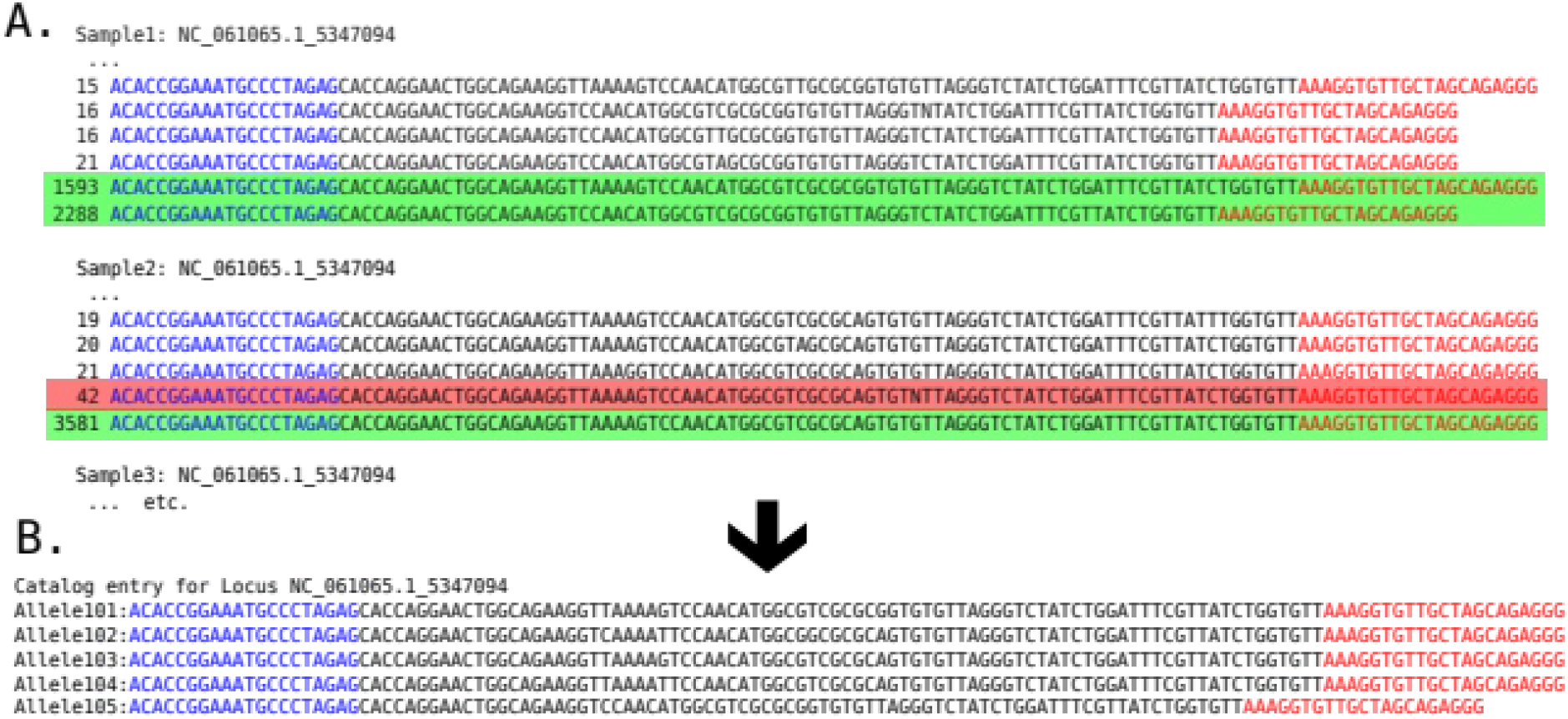
Diploid abundance model for allele discovery and catalog construction. Within each sample and locus, primer-bounded amplicon sequences are collapsed into unique sequences and their read counts are tabulated (A). Unique sequences are ranked by read abundance, and the two most abundant sequences are retained as candidate diploid alleles for each sample. Low-frequency sequences are discarded as likely sequencing artifacts (red highlight). Candidate alleles (green highlight) identified across all samples are aggregated to construct a catalog of unique haplotypes for each locus (B).

**Figure 4.**
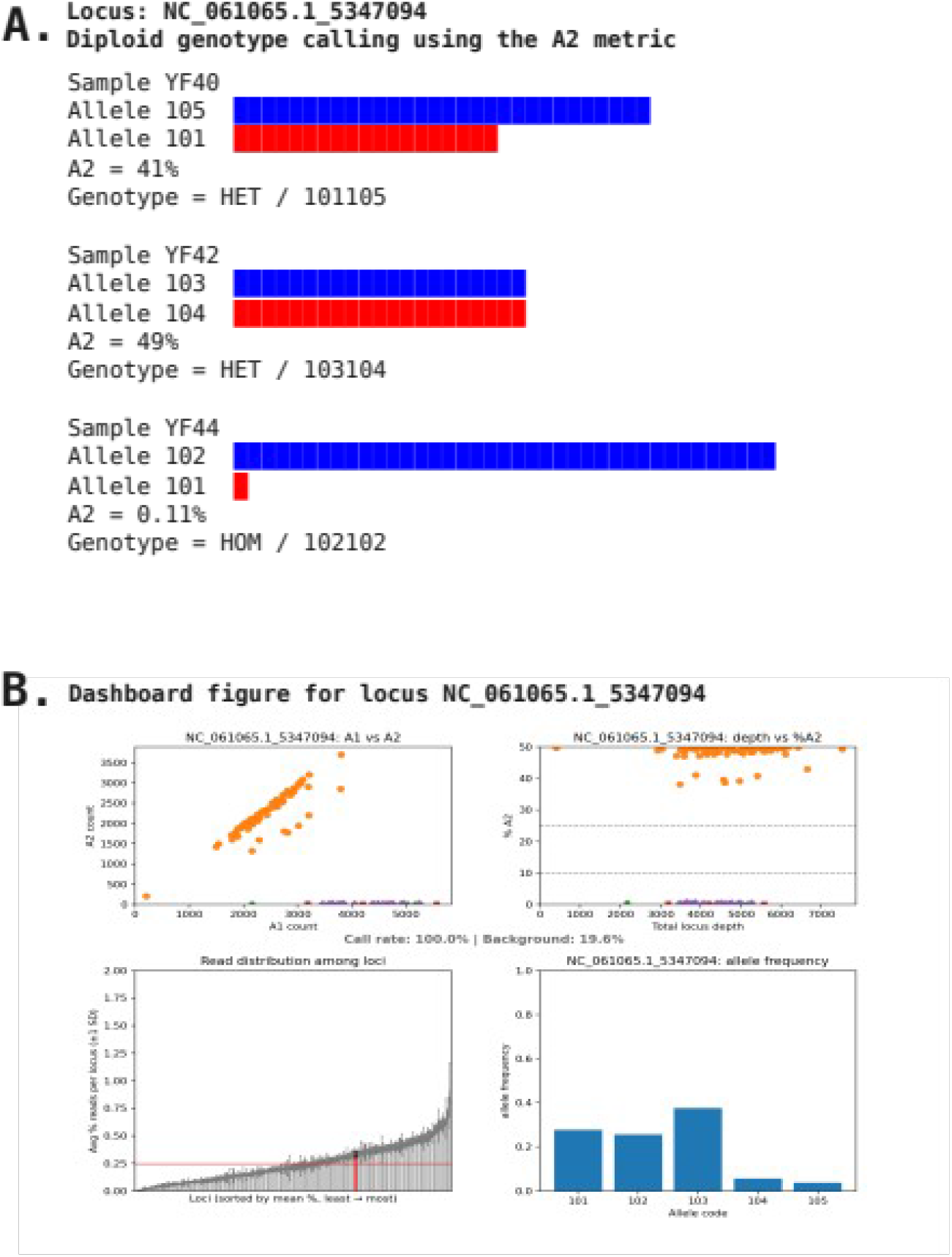
Diploid genotype inference using the A2 abundance metric and locus-level genotype summaries. (A) Diploid genotypes are inferred from read counts assigned to catalog haplotypes using the relative abundance of the second most frequent allele (A2). For each sample and locus, reads matching each catalog allele are counted and ranked by abundance. The proportion of reads corresponding to the second most abundant allele (A2) is used to distinguish heterozygous genotypes, where two alleles occur at substantial frequencies, from homozygous genotypes, where nearly all reads correspond to a single allele. Example samples from locus NC_061065.1_5347094 illustrate heterozygous and homozygous genotype calls based on allele abundance. (B) Locus-level dashboards summarize the distribution of haplotypes and diploid genotypes across samples, providing a compact representation of allele frequencies and genotype composition for each microhaplotype locus.

This alignment-free framework recovered phased haplotypes directly from GT-seq amplicon sequences without requiring read alignment or conventional variant calling (Figure 1). Because GT-seq amplicons typically span the full region containing the targeted polymorphisms, haplotype sequences could be reconstructed directly from sequencing reads following exact match genotyping (Figure 1; Figure 5).

**Figure 5.**
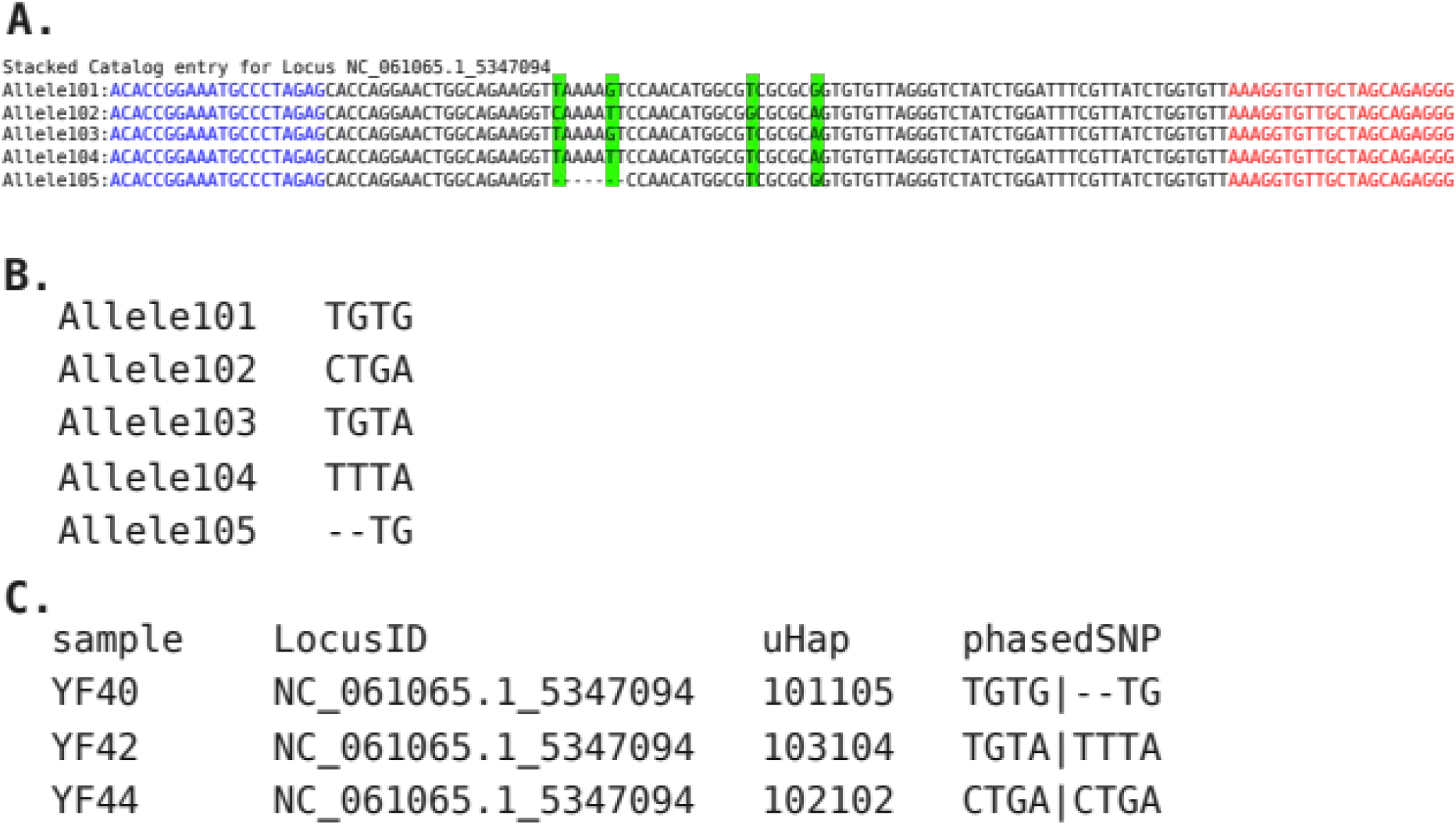
Extraction of phased microhaplotypes from catalog haplotypes. Catalog haplotypes for locus NC_061065.1_5347094 are compared to identify polymorphic positions within the amplicon, including both single nucleotide polymorphisms and insertion– deletion variants (A). Because each haplotype sequence originates from a single primer-bounded amplicon, the relative phase of polymorphisms within the locus is preserved (B). Diploid genotypes are therefore represented as combinations of phased haplotype sequences, forming multi-allelic microhaplotypes (C) suitable for downstream population genetic, parentage, and kinship analyses.

### Diploid Genotype Inference Using the A2 Metric

Diploid genotypes were inferred from allele-specific read counts using the relative abundance of the second most frequent allele (A2). For each sample and locus, reads matching catalog haplotypes were counted and ranked by abundance. The two most abundant alleles were retained as the candidate diploid genotype, and the proportion of reads corresponding to the second-most abundant allele was used to distinguish heterozygous and homozygous genotypes.

Examples from locus NC_061065.1_5347094 illustrate this approach (Figure 4A). In heterozygous individuals, two haplotypes were observed at similar frequencies, typically with A2 proportions near 0.5. In contrast, homozygous individuals exhibited extremely low A2 values, reflecting the presence of only a single dominant haplotype at the locus. This abundance-based model provided a simple and robust method for genotype inference using high-depth amplicon sequencing data.

Across the dataset, locus-level genotype summaries revealed the distribution of haplotypes and diploid genotypes across samples (Figure 4B). These locus dashboards provide a compact representation of allele frequencies and genotype composition for each microhaplotype locus. These results were consistent across sequencing platforms, with cross-platform comparisons demonstrating highly concordant genotype calls (Figure S1; Figure S2; Table S1).

### Microhaplotype Extraction and Phased SNP Representation

Catalog haplotypes were positionally compared within each locus to identify polymorphic sites, including both single nucleotide polymorphisms and short insertion–deletion variants (indels). Because each haplotype sequence represents a contiguous DNA fragment derived from a single primer-bounded amplicon, the relative phase of polymorphisms within the locus was preserved directly in the catalog haplotypes.

Locus NC_061065.1_5347094 contained multiple polymorphic positions that could be resolved into distinct haplotype sequences (Figure 5). These haplotypes represent phased combinations of SNPs and indels within the amplicon and therefore constitute microhaplotypes. Diploid genotypes at this locus were represented as pairs of haplotype sequences, producing phased microhaplotype genotypes suitable for downstream population genetic and kinship analyses.

Because haplotypes are reconstructed directly from sequencing reads rather than inferred from independent SNP calls, the resulting microhaplotypes retain the full linkage information contained within each amplicon. This representation captures the joint information from multiple polymorphic sites and can substantially increase the effective information content of loci compared to single SNP markers.

## Discussion

### Alignment-Free Microhaplotype Inference from Amplicon Sequencing Data

Targeted amplicon sequencing approaches such as GT-seq generate sequencing reads that frequently span the entire amplified region, allowing combinations of linked polymorphisms to be observed directly on individual DNA molecules. Conventional analytical pipelines typically process these data using alignment-based variant calling approaches in which sequencing reads are mapped to a reference genome and polymorphisms are identified as independent SNP markers. While effective for many applications, these workflows do not fully exploit the structure of targeted amplicon data where complete haplotypes can often be reconstructed directly from sequencing reads.

Several analytical tools have been developed to extract microhaplotypes from massively parallel sequencing data. Many of these methods rely on alignment-based workflows in which sequencing reads are first mapped to a reference genome and haplotypes are subsequently reconstructed from combinations of SNP calls. For example, the microhaplot software package identifies microhaplotypes from aligned sequencing reads and provides tools for analyzing haplotype structure across targeted loci (Baetscher et al. 2018). While these approaches are effective when reference genomes are available and read alignment is reliable, they treat haplotypes as a derived product of variant calls rather than the primary signal present in sequencing reads.

The framework described here takes the opposite approach by inferring haplotypes directly from sequencing reads prior to identifying SNP variation. Because GT-seq amplicons frequently span the full targeted region, haplotypes can be reconstructed directly from primer-bounded reads without reference alignment or statistical phasing. In this workflow, polymorphic sites are identified only after the haplotype catalog has been constructed. This inversion of the traditional analysis order— inferring haplotypes first and extracting SNP variation second—provides a simplified and computationally efficient strategy for microhaplotype genotyping from high-depth amplicon sequencing data. The high concordance observed between Element and Illumina sequencing platforms further demonstrates that this alignment-free framework is robust to differences in sequencing technology, supporting its general applicability across short-read amplicon sequencing platforms.

### Diploid Abundance Model and Catalog Construction

Central to this workflow is the diploid abundance model used to infer candidate alleles and construct locus-specific haplotype catalogs. Because GT-seq assays typically generate very high read depth per locus and Illumina and Element sequencing platforms produce predominantly high-quality reads, true allelic sequences are expected to occur at substantially higher frequencies than sequencing artifacts. Within each sample and locus, unique amplicon sequences are therefore ranked by read abundance and the two most abundant sequences are retained as candidate diploid alleles.

Aggregating these candidate alleles across samples produces a catalog of unique haplotypes for each locus that can subsequently be used for genotype inference. This abundance-based strategy provides a simple and robust approach for distinguishing true allelic sequences from sequencing artifacts in high-depth amplicon datasets.

Catalog construction assumes that each locus represents a single genomic target in a diploid organism. When multiple genomic regions are amplified by the same primer pair—such as duplicated loci or non-specific amplification events—more than two high-frequency sequences may be observed within a single sample. Under these conditions the diploid abundance model cannot unambiguously determine which sequences represent the true alleles. Rather than attempting to infer genotypes under ambiguous conditions, the algorithm intentionally fails to produce genotype calls for such loci, resulting in empty output for the affected marker. In practice this behavior serves as a useful diagnostic indicator of problematic loci, highlighting primer sets that amplify multiple genomic targets or duplicated regions of the genome. In practice, such loci are identified and excluded during panel design through in silico specificity screening and empirical optimization (as performed during development of the example dataset used here). However, application of this pipeline to existing GT-seq panels may result in missing data for some loci.

Because the current implementation assumes a maximum of two alleles per locus per individual, the method is most appropriate for single-copy loci in diploid organisms. Nevertheless, the underlying framework could be extended to accommodate more complex genotype models. For example, a modified implementation could retain the top four sequences within each sample and apply abundance-based genotype inference appropriate for tetraploid organisms.

### Microhaplotypes as Highly Informative Genetic Markers

Microhaplotypes have emerged as powerful genetic markers because they capture multiple linked polymorphisms within a single short genomic region. Unlike individual SNP markers, which are typically bi-allelic, microhaplotypes often produce multi-allelic loci with substantially higher heterozygosity and information content (Kidd et al. 2014; Pakstis et al. 2021).

Because microhaplotypes represent phased combinations of SNPs and indels observed on the same DNA fragment, each locus can contribute considerably more information to relatedness analyses than a single SNP marker. Comparative studies have demonstrated that microhaplotype panels can outperform SNP panels when distinguishing closely related individuals or resolving complex pedigree relationships (Anderson et al. 2018; Tomas et al. 2024). The multi-allelic nature of microhaplotypes increases their discriminatory power and reduces the number of loci required to achieve a given level of statistical confidence in kinship inference.

Targeted amplicon sequencing methods such as GT-seq are particularly well suited for microhaplotype genotyping because sequencing reads frequently span the entire amplified region. Consequently, many existing GT-seq panels already contain loci that can function as microhaplotypes without modification to laboratory protocols.

### Implications and Applications

The alignment-free microhaplotype pipeline described here provides a practical framework for extracting microhaplotypes directly from GT-seq amplicon sequencing data. By operating directly on primer-bounded amplicon sequences, the method avoids the need for reference alignment and variant-calling pipelines while preserving the phased structure of polymorphisms within each locus.

Because many ecological and conservation genomics studies already rely on GT-seq panels for genotyping large numbers of individuals, the ability to extract microhaplotypes from these datasets has the potential to substantially increase the information content of existing marker panels without requiring changes to laboratory protocols. In this sense, the alignment-free framework described here provides a practical path for converting existing SNP-based GT-seq panels into multi-allelic microhaplotype marker systems.

More broadly, this work demonstrates how haplotype-centric analytical approaches can simplify the interpretation of targeted sequencing data. By treating haplotypes as the primary signal present in amplicon reads and identifying polymorphic sites only after haplotypes have been inferred, the alignment-free framework provides an efficient strategy for microhaplotype genotyping from high-depth amplicon sequencing experiments.

## Software Availability

The analysis pipeline described in this study is implemented as a Python script available through the GTseq GitHub repository:

https://github.com/GTseq/gtseq_microhap

The repository contains the source code, documentation, and example usage instructions for performing alignment-free microhaplotype genotyping from GT-seq paired-end sequencing data.

## Data Availability

All sequencing data and example datasets are available at Zenodo: https://doi.org/10.5281/zenodo.19069551. The analysis pipeline is available at https://github.com/GTseq/gtseq_microhap.

Cross-platform genotype concordance analyses, including sample-level concordance metrics (Table S1) and visualization of locus-level and pairwise genotype agreement (Figures S1–S2), are provided as supplementary materials.

## Author Contributions

Identification and selection of microhaplotype markers for kinship analysis in delta smelt were carried out by SKB and AJF. The analytical framework, software pipeline, and manuscript were developed by NRC. Software testing and manuscript review and revision were contributed by ARC, SKB, and AJF. GT-seq library preparation and laboratory operations were performed and overseen by ARC.

