## Supplementary material for "Alignment-Free Microhaplotype Genotyping for GT-seq (Genotyping-in-Thousands by Sequencing) Using a Diploid Abundance Model": Figure S1

Figure S1: Locus-by-sample genotype concordance between Element and Illumina sequencing platforms. Green indicates matching genotype calls, red indicates mismatches, and yellow/orange indicate comparisons involving no-calls.

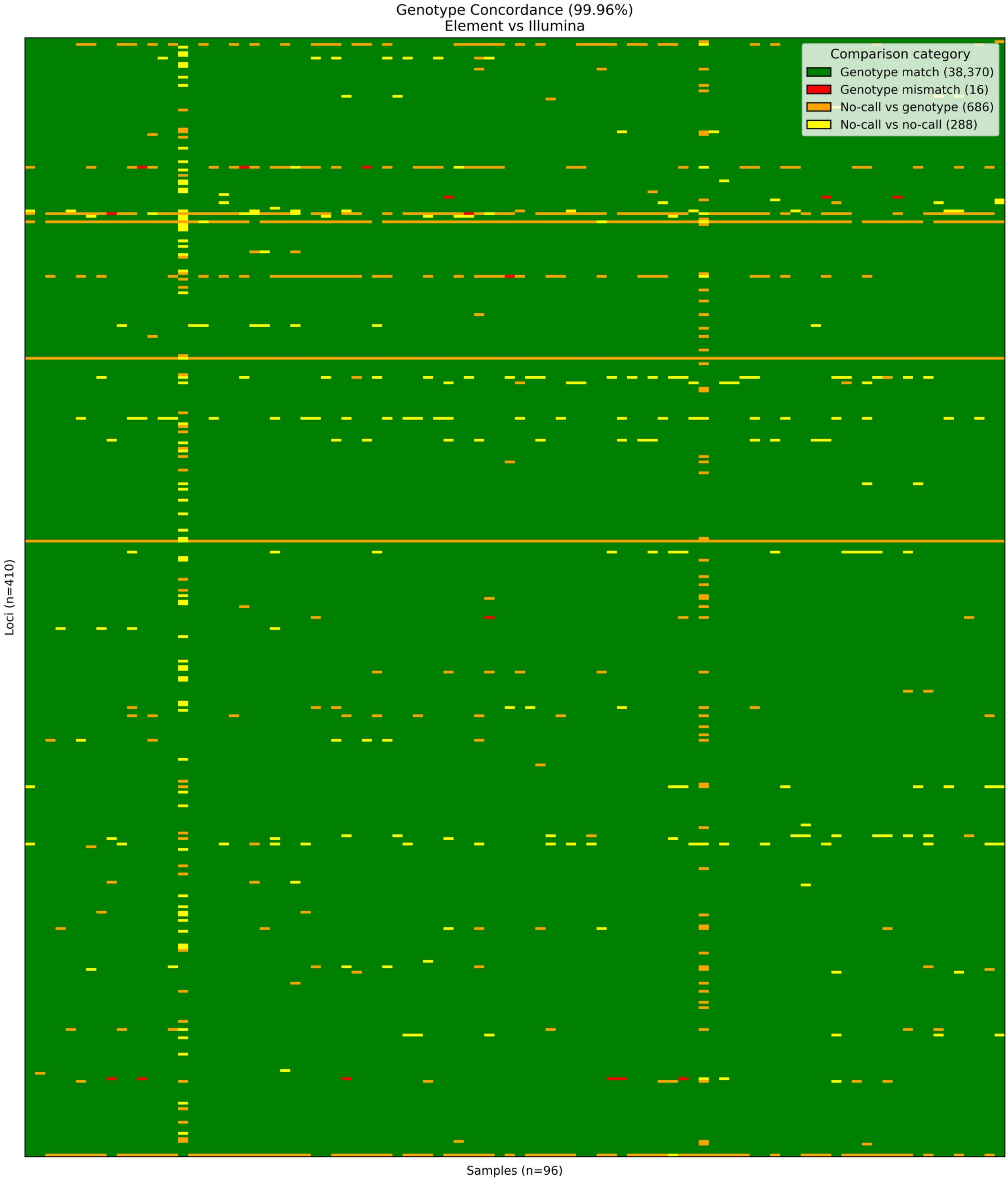
