## Supplementary figures and images for "Alignment-Free Microhaplotype Genotyping for GT-seq (Genotyping-in-Thousands by Sequencing) Using a Diploid Abundance Model"

### Figure S2

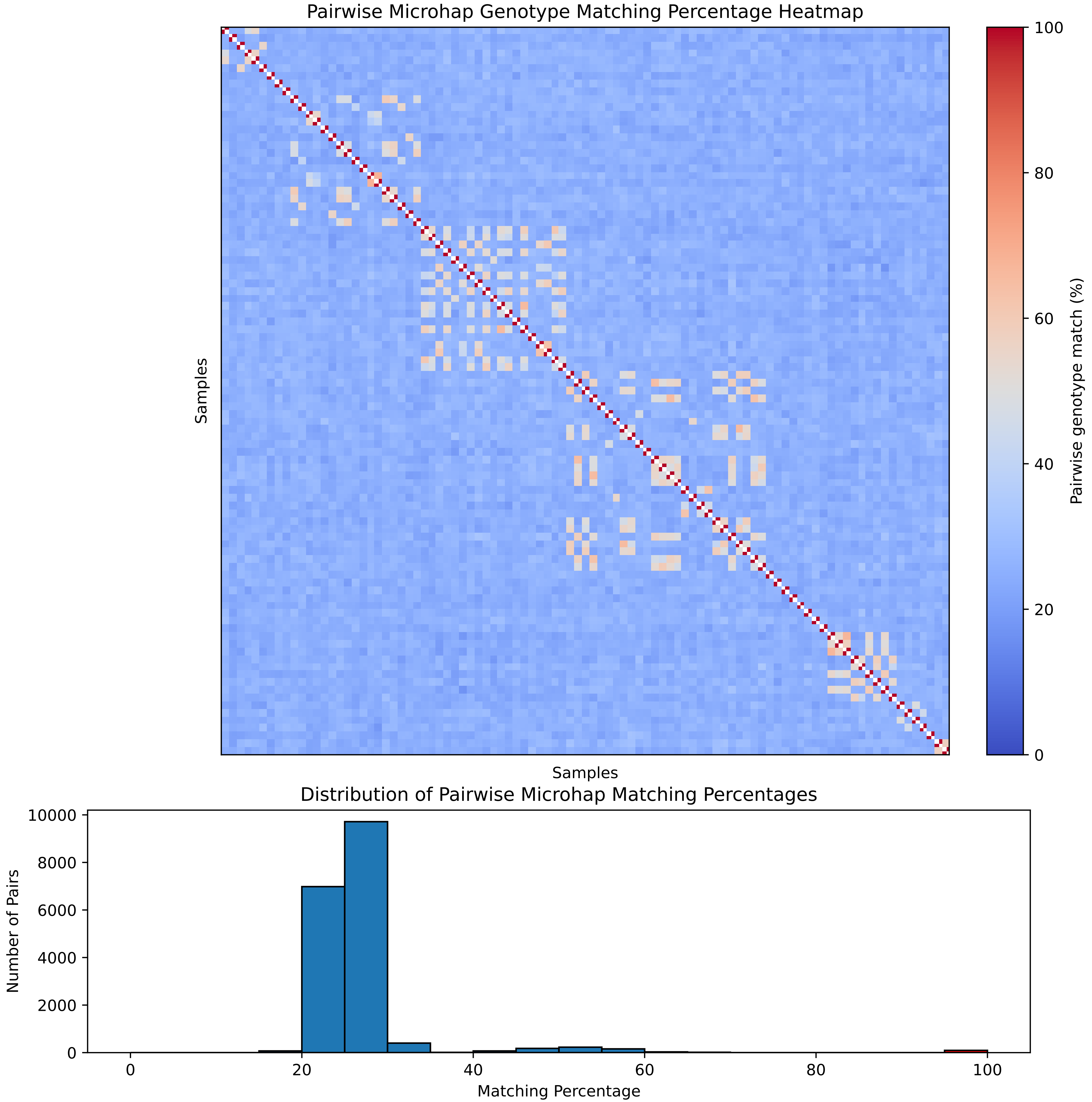
