## Supplementary material for "Alignment-Free Microhaplotype Genotyping for GT-seq (Genotyping-in-Thousands by Sequencing) Using a Diploid Abundance Model": Table S1

### Table S1. Sample-level genotype concordance between Element and Illumina sequencing runs

Percent match is calculated across loci successfully genotyped in both datasets. All corresponding sample pairs exhibit >99.5% concordance.

| Element Sample | Illumina Sample | % Match | %GT Element | %GT Illumina |
|---|---|---|---|---|
| Element_OC31_1 | Illumina_OC31_1 | 100.00% | 99.0% | 98.3% |
| Element_OC32_1 | Illumina_OC32_1 | 100.00% | 99.8% | 99.5% |
| Element_OC33_1 | Illumina_OC33_1 | 100.00% | 100.0% | 98.3% |
| Element_OC37_2 | Illumina_OC37_2 | 100.00% | 99.8% | 98.3% |
| Element_OC38_2 | Illumina_OC38_2 | 100.00% | 99.8% | 98.3% |
| Element_OC42_2 | Illumina_OC42_2 | 100.00% | 99.5% | 97.6% |
| Element_OF02_2 | Illumina_OF02_2 | 100.00% | 99.5% | 97.6% |
| Element_OG07_1 | Illumina_OG07_1 | 100.00% | 99.5% | 97.8% |
| Element_YA88_2 | Illumina_YA88_2 | 99.50% | 99.5% | 98.5% |
| Element_YB69_2 | Illumina_YB69_2 | 100.00% | 99.5% | 97.8% |
| Element_YB70_1 | Illumina_YB70_1 | 100.00% | 99.0% | 97.3% |
| Element_YB71_1 | Illumina_YB71_1 | 99.51% | 99.8% | 98.8% |
| Element_YB72_1 | Illumina_YB72_1 | 100.00% | 99.5% | 97.6% |
| Element_YB74_2 | Illumina_YB74_2 | 100.00% | 99.5% | 98.3% |
| Element_YB77_2 | Illumina_YB77_2 | 100.00% | 99.5% | 97.6% |
| Element_YB79_2 | Illumina_YB79_2 | 100.00% | 99.8% | 98.5% |
| Element_YB80_2 | Illumina_YB80_2 | 100.00% | 99.5% | 98.0% |
| Element_YB81_2 | Illumina_YB81_2 | 100.00% | 99.8% | 98.3% |
| Element_YB82_2 | Illumina_YB82_2 | 100.00% | 99.3% | 97.8% |
| Element_YB85_2 | Illumina_YB85_2 | 100.00% | 100.0% | 98.0% |
| Element_YB86_2 | Illumina_YB86_2 | 99.75% | 99.5% | 98.0% |
| Element_YB87_2 | Illumina_YB87_2 | 100.00% | 99.0% | 97.6% |
| Element_YB88_2 | Illumina_YB88_2 | 100.00% | 99.5% | 98.0% |
| Element_YB89_2 | Illumina_YB89_2 | 100.00% | 98.5% | 96.8% |
| Element_YB90_2 | Illumina_YB90_2 | 100.00% | 99.8% | 97.8% |
| Element_YB91_2 | Illumina_YB91_2 | 100.00% | 98.5% | 97.3% |
| Element_YF40_1 | Illumina_YF40_1 | 100.00% | 99.5% | 97.8% |
| Element_YF41_1 | Illumina_YF41_1 | 100.00% | 99.0% | 97.6% |
| Element_YF42_1 | Illumina_YF42_1 | 100.00% | 99.5% | 98.0% |

| Element Sample | Illumina Sample | % Match | %GT Element | %GT Illumina |
|---|---|---|---|---|
| Element_YF43_1 | Illumina_YF43_1 | 100.00% | 99.3% | 97.3% |
| Element_YF44_2 | Illumina_YF44_2 | 99.75% | 98.8% | 97.1% |
| Element_YF45_2 | Illumina_YF45_2 | 100.00% | 100.0% | 97.8% |
| Element_YF46_2 | Illumina_YF46_2 | 99.75% | 99.5% | 98.3% |
| Element_YF48_2 | Illumina_YF48_2 | 100.00% | 99.3% | 97.8% |
| Element_YF49_2 | Illumina_YF49_2 | 100.00% | 99.0% | 97.1% |
| Element_YF50_2 | Illumina_YF50_2 | 100.00% | 99.5% | 98.8% |
| Element_YF53_2 | Illumina_YF53_2 | 100.00% | 99.0% | 97.8% |
| Element_YF54_2 | Illumina_YF54_2 | 100.00% | 99.5% | 97.8% |
| Element_YF55_2 | Illumina_YF55_2 | 100.00% | 99.3% | 97.3% |
| Element_YF56_2 | Illumina_YF56_2 | 100.00% | 99.5% | 98.3% |
| Element_YF57_2 | Illumina_YF57_2 | 99.75% | 98.8% | 97.3% |
| Element_YF58_2 | Illumina_YF58_2 | 100.00% | 99.5% | 97.6% |
| Element_YF59_2 | Illumina_YF59_2 | 99.75% | 99.5% | 98.0% |
| Element_YF63_1 | Illumina_YF63_1 | 100.00% | 98.5% | 96.8% |
| Element_YF80_2 | Illumina_YF80_2 | 99.75% | 99.3% | 97.1% |
| Element_YG31_1 | Illumina_YG31_1 | 100.00% | 100.0% | 98.3% |
| Element_YG33_1 | Illumina_YG33_1 | 99.75% | 99.5% | 98.0% |
| Element_YG34_1 | Illumina_YG34_1 | 100.00% | 99.8% | 97.8% |
| Element_YG35_1 | Illumina_YG35_1 | 100.00% | 99.5% | 98.5% |
| Element_YG37_1 | Illumina_YG37_1 | 100.00% | 99.3% | 97.8% |
| Element_YG38_1 | Illumina_YG38_1 | 100.00% | 99.5% | 97.6% |
| Element_YG39_1 | Illumina_YG39_1 | 100.00% | 99.8% | 98.0% |
| Element_YG40_1 | Illumina_YG40_1 | 100.00% | 99.3% | 97.8% |
| Element_YG41_1 | Illumina_YG41_1 | 100.00% | 99.8% | 98.3% |
| Element_YG42_1 | Illumina_YG42_1 | 100.00% | 99.8% | 98.3% |
| Element_YG43_2 | Illumina_YG43_2 | 100.00% | 99.5% | 97.6% |
| Element_YG44_2 | Illumina_YG44_2 | 99.75% | 99.5% | 98.3% |
| Element_YG45_2 | Illumina_YG45_2 | 99.75% | 99.0% | 98.0% |
| Element_YG46_2 | Illumina_YG46_2 | 100.00% | 99.8% | 98.3% |
| Element_YG47_2 | Illumina_YG47_2 | 100.00% | 99.8% | 98.0% |
| Element_YG50_2 | Illumina_YG50_2 | 100.00% | 99.3% | 97.6% |
| Element_YG51_2 | Illumina_YG51_2 | 100.00% | 99.5% | 97.6% |
| Element_YG52_2 | Illumina_YG52_2 | 100.00% | 98.5% | 97.1% |

| Element Sample | Illumina Sample | % Match | %GT Element | %GT Illumina |
|---|---|---|---|---|
| Element_YG53_2 | Illumina_YG53_2 | 99.75% | 99.3% | 97.6% |
| Element_YG54_2 | Illumina_YG54_2 | 100.00% | 99.0% | 98.0% |
| Element_YG55_2 | Illumina_YG55_2 | 100.00% | 98.0% | 84.1% |
| Element_YG57_2 | Illumina_YG57_2 | 100.00% | 99.8% | 98.5% |
| Element_YG58_1 | Illumina_YG58_1 | 100.00% | 99.0% | 97.8% |
| Element_YG61_1 | Illumina_YG61_1 | 100.00% | 99.8% | 98.5% |
| Element_YG62_1 | Illumina_YG62_1 | 100.00% | 99.5% | 98.5% |
| Element_YG64_1 | Illumina_YG64_1 | 100.00% | 99.3% | 97.3% |
| Element_YH07_1 | Illumina_YH07_1 | 100.00% | 99.8% | 98.5% |
| Element_YI07_2 | Illumina_YI07_2 | 100.00% | 99.3% | 98.0% |
| Element_YK00_1 | Illumina_YK00_1 | 100.00% | 99.5% | 98.0% |
| Element_YK01_2 | Illumina_YK01_2 | 100.00% | 99.5% | 98.5% |
| Element_YL06_1 | Illumina_YL06_1 | 100.00% | 99.3% | 98.0% |
| Element_YN17_2 | Illumina_YN17_2 | 100.00% | 99.5% | 98.5% |
| Element_YO11_1 | Illumina_YO11_1 | 99.75% | 99.5% | 98.0% |
| Element_YO12_2 | Illumina_YO12_2 | 100.00% | 98.3% | 97.1% |
| Element_YR30_1 | Illumina_YR30_1 | 100.00% | 99.5% | 98.0% |
| Element_YR33_2 | Illumina_YR33_2 | 100.00% | 99.8% | 98.5% |
| Element_YR34_2 | Illumina_YR34_2 | 100.00% | 99.0% | 97.3% |
| Element_YR35_2 | Illumina_YR35_2 | 100.00% | 99.3% | 98.0% |
| Element_YR36_2 | Illumina_YR36_2 | 100.00% | 99.5% | 98.3% |
| Element_YR37_2 | Illumina_YR37_2 | 99.75% | 100.0% | 98.8% |
| Element_YR38_2 | Illumina_YR38_2 | 100.00% | 99.3% | 97.8% |
| Element_YR39_2 | Illumina_YR39_2 | 100.00% | 99.5% | 98.5% |
| Element_YR40_2 | Illumina_YR40_2 | 100.00% | 99.0% | 98.0% |
| Element_YT27_1 | Illumina_YT27_1 | 100.00% | 99.3% | 98.0% |
| Element_YT28_1 | Illumina_YT28_1 | 100.00% | 99.3% | 98.3% |
| Element_YT30_1 | Illumina_YT30_1 | 100.00% | 99.3% | 97.8% |
| Element_YT32_1 | Illumina_YT32_1 | 100.00% | 100.0% | 98.5% |
| Element_YX28_1 | Illumina_YX28_1 | 100.00% | 99.8% | 98.5% |
| Element_YX30_2 | Illumina_YX30_2 | 100.00% | 99.5% | 97.6% |
| Element_YX32_2 | Illumina_YX32_2 | 100.00% | 98.8% | 97.8% |
